# Binding affinity and conformational preferences influence kinetic stability of short oligonucleotides on carbon nanotubes

**DOI:** 10.1101/2020.02.08.939918

**Authors:** Ali A. Alizadehmojarad, Xingcheng Zhou, Abraham G. Beyene, Kevin Chacon, Younghun Sung, Markita P. Landry, Lela Vuković

## Abstract

DNA-wrapped single walled carbon nanotubes (SWNTs) have found a widespread use in a variety of nanotechnology applications. Yet, the relationship between structural conformation, binding affinity and kinetic stability of these polymers on SWNTs remains poorly understood. Here, we used molecular dynamics (MD) simulations and experiments to explore this relationship for short oligonucleotides adsorbed on SWNTs. First, using classical MD simulations of oligonucleotide-(9,4)-SWNT hybrid complexes, we explored the relationship between ssDNA and ssRNA surface conformation and sequence chemistry. We screened the conformation of 36 sequences of short ssDNA and ssRNA polymers on (9,4) SWNT, where the contour lengths were selected so the polymers can, to a first approximation, wrap once around the SWNT circumference. From these screens, we identified structural motifs that we broadly classified into “rings” and “non-rings.” Then, several sequences were selected for detailed investigations. We used temperature replica exchange MD calculations to compute two-dimensional free energy landscapes characterizing the conformations of select sequences. “Ring” conformations seemed to be driven primarily by sequence chemistry. Specifically, strong (n,n+2) nucleotide interactions and the ability of the polymer to form compact structures, as for example, through sharp bends in the nucleotide backbone, correlated with ring-forming propensity. However, ring-formation probability was found to be uncorrelated with free energy of oligonucleotide binding to SWNTs (ΔG_bind_). Conformational analyses of oligonucleotides, computed free energy of binding of oligonucleotides to SWNTs, and experimentally determined kinetic stability measurements show that ΔG_bind_ is the primary correlate for kinetic stability. The probability of the sequence to adopt a compact, ring-like conformation is shown to play a secondary role that still contributes measurably to kinetic stability. For example, sequences that form stable compact rings (C-rich sequences) could compensate for their relatively lower ΔG_bind_ and exhibit kinetic stability, while sequences with strong ΔG_bind_ (such as (TG)_3_(GT)_3_) were found to be kinetically stable despite their low ring formation propensity. We conclude that the stability of adsorbed oligonucleotides is primarily driven by its free energy of binding and that if ring-like structural motifs form, they would contribute positively to stability.

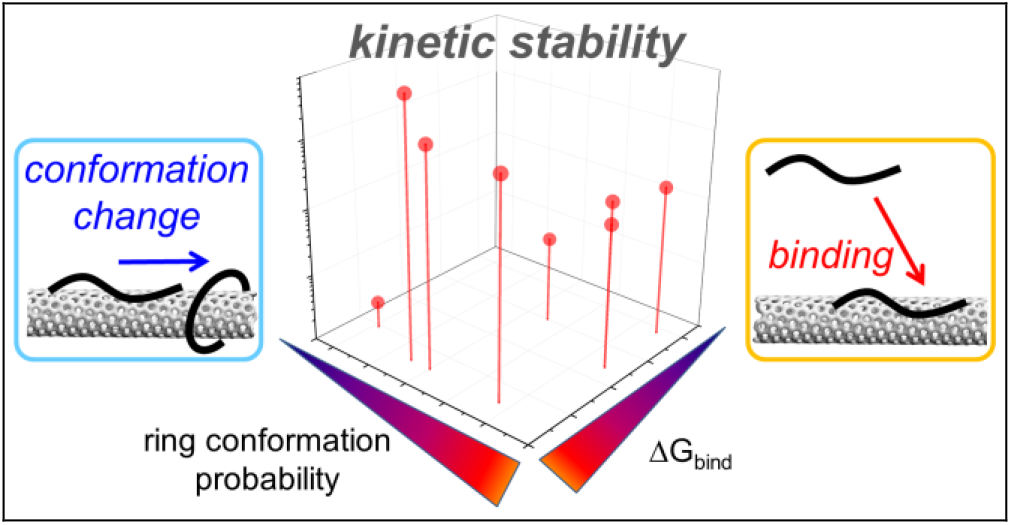

## 1. INTRODUCTION

Single wall carbon nanotubes (SWNTs) comprise a key component of many new nanotechnology applications for sensing, biological imaging, electronics, and gene delivery, among others^1–11^. Noncovalent polymer adsorption is a widely used method to confer and optimize desired functionalities to SWNTs, while solubilizing them in aqueous environments. A variety of polymers, which when adsorbed form a “corona phase” on the surface of SWNT, have been used for SWNT functionalization, including nucleic acids, peptides, peptoids, lipids, surfactants, and others^12–20^. Among those, nucleic acid functionalized SWNTs in particular were found to have a number of technologically useful and important applications^5, 10, 21–24^. This variety of polymers and the ease of functionalization through noncovalent adsorption provide a great opportunity for rational design of SWNT conjugates for specific applications. However, rational design methodologies for corona phase engineering would benefit from a systematic body of knowledge about how individual polymer properties, such as polymer length, sequence, and chemistry, relate to structural and kinetic properties of the resulting corona phase. As polymer corona cannot be directly visualized with atomic resolution in experiments, important characteristics of the corona phase, such as structural arrangement of polymers, corona thickness, dynamics of assembled polymers on different timescales, kinetic stability of polymer corona, and interactions with other molecules present in the suspension, have to be inferred from experiments with limited resolution^25–29^.

An alternative and a powerful way to characterize the corona phase includes computational modeling approaches, such as classical molecular dynamics (MD) simulations, which enable atomic resolution on the timescale of hundreds of nanoseconds^1, 2, 29–32^. Yet, as reorganization of long polymers on SWNT surfaces occurs on a timescale of several nanoseconds, determining most probable equilibrium distributions of polymers in the corona phase necessitates enhanced sampling approaches, such as temperature replica exchange (T-REMD) MD simulations^33^. Complete free energy landscapes and equilibrium distributions of select representative polymers, such as single DNA sequences, on SWNT surface were determined in prior studies^2, 30, 34^. While the number of systems sampled in simulations has typically been small, limited to several sequences at most, investigating a larger number of systems recently became more computationally feasible.

Generating extended sets of data from systematically sampled atomistic resolution simulations could open attractive routes for developing predictive frameworks for corona phase design, which could overcome the present severe limits of *ab initio* predictions of polymers with specific functions, such as SWNT or analyte recognition. Several recent experimental studies already address the lack of big data in the polymer-functionalized SWNT field in general, utilizing high throughput approaches to discover polymers (nucleic acid sequences) which can recognize specific chiralities of SWNTs or sense particular analytes, such as serotonin^35, 36^. In the present paper, we select an extended set of related nucleic acid polymers, which vary in sequence but have similar short lengths, and combine extended atomistic-resolution computational screening (MD, T-REMD simulations) with lower-resolution experimental screening (kinetic stability surfactant displacement experiments) to relate and generalize the relationships between structural conformation, binding affinity and kinetic stability of short oligonucleotides adsorbed on SWNTs.

## 2. RESULTS AND DISCUSSION

### 2.1. Screening conformations of circumference-length nucleic acid polymers on (9,4) SWNTs

In a previous study, we reported results from molecular dynamics (MD) study showing that the oligonucleotide sequence, (GT)_6_, formed semi-ordered, ring-like structures when adsorbed on (9,4)-SWNT but did not do so when adsorbed on the (6,5)-SWNT. Numerous studies have previously investigated surface conformations of oligonucleotides adsorbed on SWNTs and identified helical structures as the general predominant form. The ring-like structural motifs were unique to our study and suggested that short oligonucleotides adsorbed on bigger diameter SWNT may form ordered ring like structures that render unique photophysical and functional properties to oligonucleotide-SWNT hybrid structure. Here, we extend this work to explore relationships between sequence chemistry, binding affinity and surface free energy landscapes to gain better understanding for ring forming sequences and identify physical rules that underpin their formation. Towards this end, we screened the conformations of 36 short single stranded nucleic acids on (9,4) SWNTs through single-trajectory equilibrium MD simulations. The sequences screened included 10-, 11-, and 12-nucleotide (nt) long ssDNAs and ssRNAs, whose lengths, to a first approximation, matched the circumference of the (9,4)-SWNTs (Fig. 1a and Table S2). These nucleic acid strands were initially equilibrated with left-handed helix configurations around (9,4) SWNTs in 0.1 M NaCl aqueous solution. After minimization and brief (2 ns) equilibration of the ions around the restrained nucleic acids and SWNTs, the nucleic acids were released from restraints and equilibrated for an additional 80 ns.

**Figure 1.**
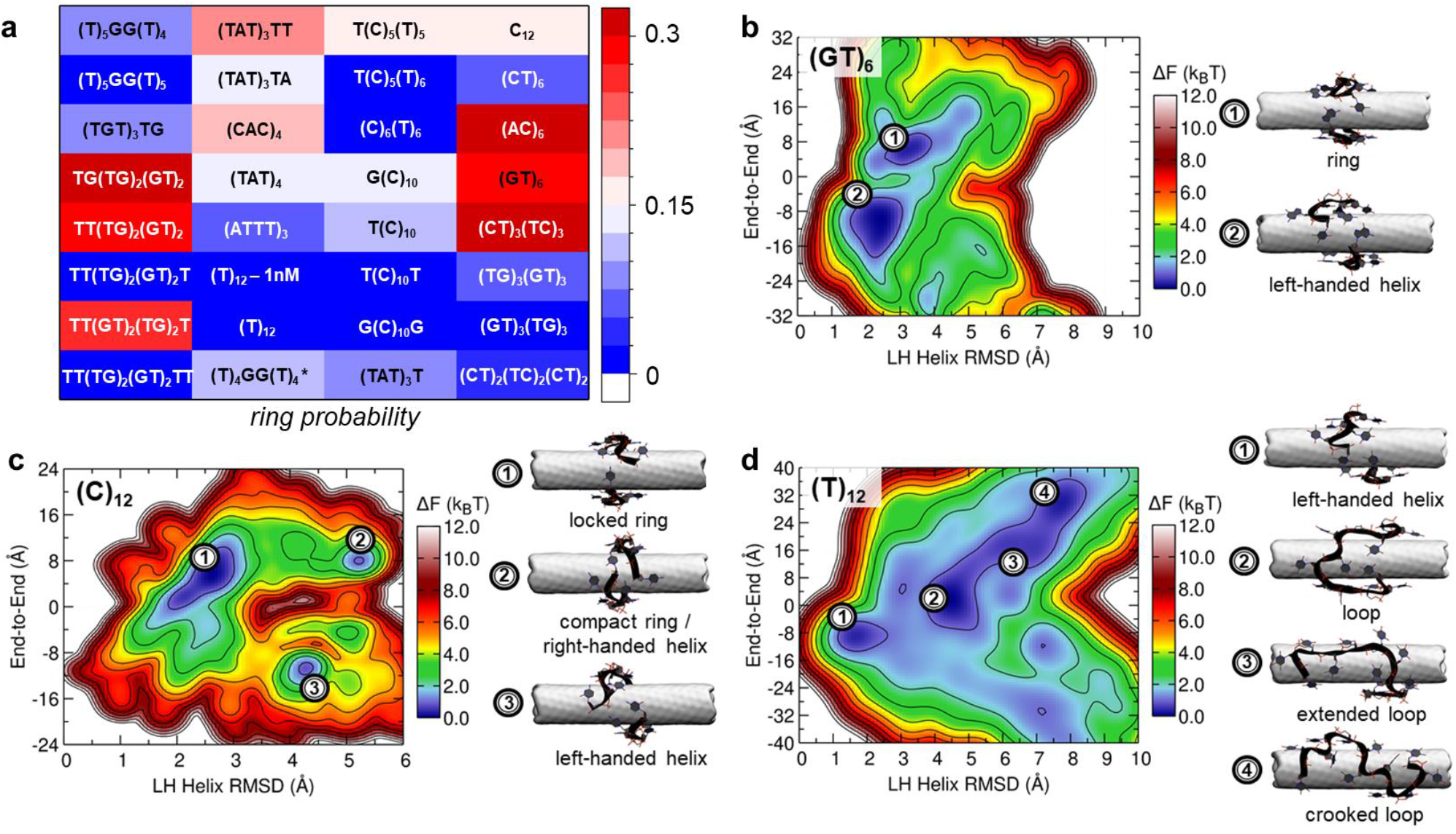
Single stranded nucleic acid conformations on (9,4) SWNT at 300 K. a. Screening of 32 sequences for ring conformation probabilities on (9,4) SWNTs in short single trajectory MD simulations at 300 K. b. Free energy landscape of (GT)_6_ on (9,4) SWNT in T-REMD calculation at 300 K. (GT)_6_-SWNT has two free energy minima, corresponding to ring (1) and left-handed (2) helix conformations. b. Free energy landscape of C_12_ on (9,4) SWNT in T-REMD calculation at 300 K. C_12_-SWNT has free energy minima corresponding to locked ring (1), compact ring/right-handed helix (2), and left-handed helix conformations (3). c) Free energy landscape of T_12_ on (9,4) SWNT in T-REMD calculation at 300 K. T_12_-SWNT has four free energy minima corresponding to left-handed helix (1), loop (2), and extended (3) and crooked loop (4) conformations.

Heatmap in Figure 1a and Table S2 summarize the results of the screening performed and quantify the probability of ring formation. While ~31% of the sequences had a strong probability (>0.15) to form rings around (9,4) SWNTs, ~40% of the sequences had a low probability (<0.05) or were not observed to form ring conformations. The remaining 28% of the sequences screened showed intermediate probability to form ring conformations around (9,4) SWNTs. Multiple sequences adopted ring conformations, including the previously examined (GT)_6_^2^. Several independent MD simulations were performed for several sequences, and in one of them C_12_ ssDNA formed the most stable rings that exhibited a perpendicular arrangement with respect to the SWNT axis. A variety of ring-forming conformations were observed, where the rings were sometimes either perpendicular to or tilted with respect to the SWNT axis. The rings formed via hydrogen bonding between 3’- and 5’-end nucleotides, with hydrogen bonds forming between polar and hydrogen atoms on either the backbones, sugar rings, or bases of these nucleotides. Many different hydrogen bonds are observed to stabilize the observed rings on SWNTs, unlike the well-defined covalent bonding in circular nucleic acids^37^. Some sequences, including T_12_, did not form any ring conformations in the screening simulations. To further confirm the existence of ring- versus helix-forming structures on SWNT, we performed high resolution TEM microscopy of two previously reported ring- and helix-forming structures, short (GT)_6_ and long (GT)_15_ molecules, respectively, on SWNT surfaces. TEM images are suggestive of the predicted ring and helix conformations (Fig. S1), suggesting that sequences identified herein as putatively ring-forming can be classified as such with our combined MD and experimental approach.

Several related short ssRNA sequences were also screened for ring formation (Table S2). They displayed different behavior from ssDNA sequences. Namely, RNA sequences formed either no ring structures ((CU)_5_) or had a large probability to form transient rings that are significantly more tilted with respect to SWNT axes than DNA sequences, but whose e 3’- and 5’-end nucleotides bond transiently via hydrogen bonding ((CU)_5_UU, (CU)_6_, and (CU)_5_C), (Table S2). Transience of RNA ring structures in comparison to DNA ring structures was noted in three dimensional scatter plots of end-to-end distances and RMSD values of all structures with respect to the reference structure for each time frame (Methods). For RNA trajectories, the points representing rings in the scatter plots did not form well separated clusters, whereas for most of the DNA trajectories the points corresponding to ring structures formed clusters. The presence of the ribose sugar oxygen on RNA was previously correlated to the diminished stability of ssRNAs on SWNTs in comparison to ssDNAs^38^. A similar effect could potentially be contributing to the smaller stability of ssRNA rings in comparison to ssDNA rings.

### 2.2. Free energy landscapes of circumference-matching oligonucleotides on (9,4) SWNTs

Based on the screening reported in Fig. 1a, multiple nucleic acid sequences with different ring-formation behavior in short single trajectory MD simulations were selected for detailed investigation based on better conformational sampling^2, 34^. Namely, (GT)_6_, C_12_, (CT)_6_, (CAC)_4_, (TG)_3_(GT)_3_, and (CT)_3_(TC)_3_ DNA sequences (weak to strongly ring-forming in single trajectory MD simulations), T_12_ and (GT)_3_(TG)_3_ (not ring forming in single trajectory MD simulations) DNA sequences, and (CU)_6_ and (CU)_5_UU (both ring-forming) RNA sequences were examined in T-REMD calculations. REMD simulations permit detailed explorations of free energy landscapes that may not be fully sampled in single trajectory MD simulations^2, 34^. The free energy landscapes and the structures representative of the free energy minima for these sequences obtained from T-REMD calculations at T = 300 K are shown in Figs. 1b-d, S2 and S3. All the studied sequences showed multiple free energy minima, corresponding to different representative surface structures. The sequences with significant free energy minima basins that represent ring conformations include (GT)_6_ (~40% of the ensemble), C_12_ (~90% of the ensemble), (CT)_3_(TC)_3_ (~90% of the ensemble, localized and ordered), and (GT)_3_(TG)_3_ (~75% of the ensemble). Interestingly, single-trajectory MD simulations did not identify (GT)_3_(TG)_3_ as a ring-forming sequence. The other free energy minima these sequences exhibited corresponded to left-handed and right-handed helices.

Three related DNA and RNA sequences ((CT)_6_ DNA, (CU)_6_ RNA, and (CU)_5_UU RNA sequences) were also examined (Fig. S2). These related sequences all show high ring forming propensity and have free energy landscapes with similar features in which the free energy minima exhibit elongated diagonal basins. The similar features of the free energy minima for both DNA and RNA sequences indicate that both base identity and order strongly influence favored conformations of oligonucleotides.

Two sequences, T_12_ and (TG)_3_(GT)_3_, showed no or weak ring minima, but (TG)_3_(GT)_3_ had a strong minimum associated with compact left-handed helix structures. The latter sequence, (TG)_3_(GT)_3_, was classified as a ring forming sequence in the single-trajectory MD screening calculations. The two specified sequences were also the only sequences that exhibited flexible and disordered loop structures, which did not wrap SWNTs around the circumference.

The observation that some sequences did not exhibit rings in screening simulations, but did so in T-REMD calculations that have improved conformational sampling, and vice versa, as especially observed for (GT)_3_(TG)_3_ and (TG)_3_(GT)_3_, indicates that 12-nt ssDNA molecules do not fully explore their conformational space on SWNTs in single trajectory equilibrium MD simulations on the timescale of ~100 ns.

### 2.3. Ring conformations form compact structures

Since many free energy minimum structural ensembles obtained in T-REMD calculations contained ring structures (Figs. 1b-d, S2 and S3), these structures were analyzed to identify structural properties that could characterize them. Visually, two factors seem to be correlated with ring conformations. First, ring conformations often exhibit kinks in the backbone, as shown schematically in the inset of Fig. 2a. With such backbone, the ring structure appears to be more compact than other types of conformations observed, such as helical structures (intermediately compact) and loop (least compact). In ring structures, hydrogen bonding (such as between bases or backbones of (n,n+2) nucleotides, shown schematically in the inset of Fig. 2a) and bound ions appear to stabilize the compact conformations.

**Figure 2.**
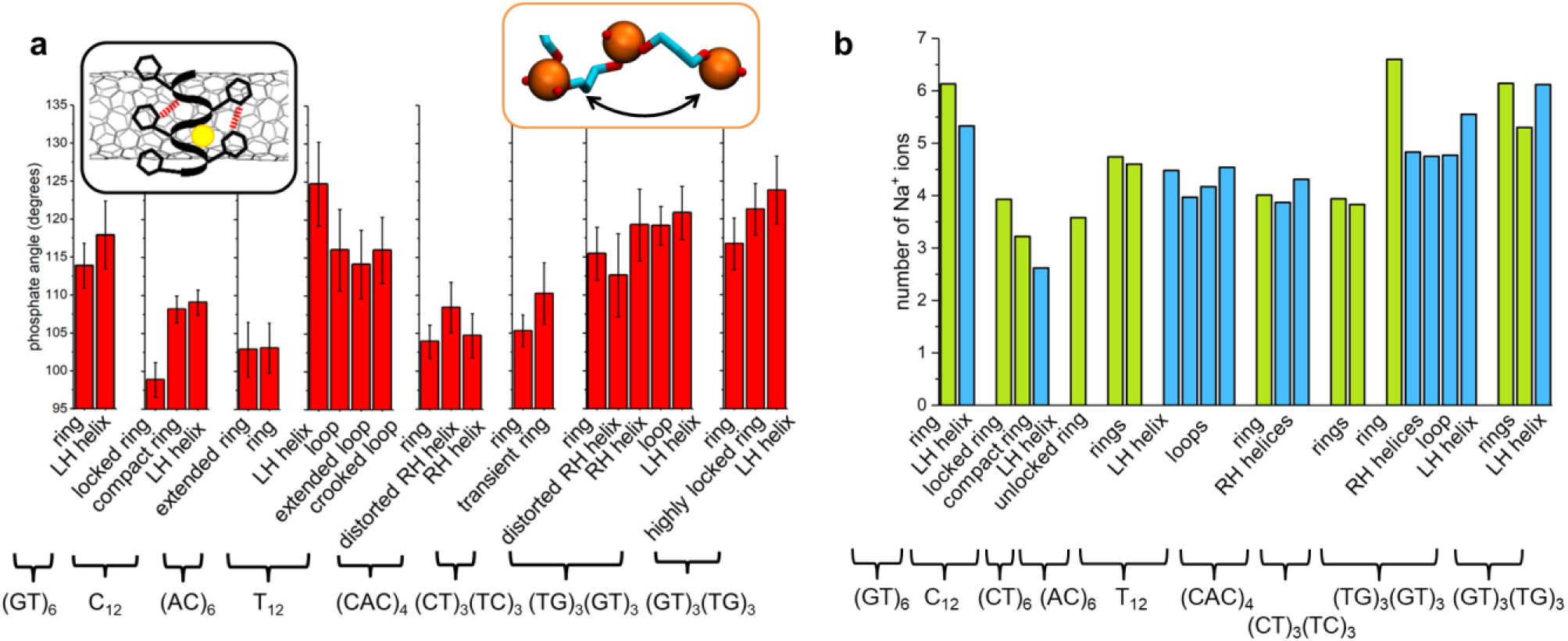
Analysis of conformational ensembles. a. Phosphate angles (orange inset) in major nucleic acids conformations, averaged over all P-atom triplets and over all the structures within the specified free energy minimum basins, determined from T-REMD calculations at 300 K. Black inset: a scheme of the ring conformation, exhibiting zig-zag structure of the backbone, which is stabilized via (n,n+2) hydrogen bonding and bound ions. b. Average number of sodium ions within 4 Å of nucleic acids examined, averaged over all the structures in free energy minimum basins, determined from T-REMD simulations at 300 K. Bars corresponding to ring conformations are shown in green, while bars corresponding to other conformations are shown in blue. Abbreviations LH and RH refer to left-handed and right-handed helices. The table with the values and full names of conformations are given in Table S3.

To quantify if ring conformations on average have a more compact backbone with zig-zag structure (kinks) in it, we calculated the angles between phosphorus (P) atoms of three neighboring phosphate groups, averaged over all triplets of P-atoms on neighboring nucleotides and over all the structures in the specified free energy minimum basins. The average angles we obtained, whose values range from ~95° to ~125°, are shown in Fig. 2a. For most of the sequences examined, ring conformations exhibit the smallest angles among the major conformations observed, and therefore the sharpest turns (kinks) in the backbone.

Ring conformations also appeared to be more compact than other conformations due to the presence of ions and potentially hydrogen bonds between (n, n+2) nucleotides. The nucleic acids occasionally form pockets in which ions are stabilized for long times, as reported in our previous study^2^. The average numbers of ions at close distance (within 4 Å) from nucleic acids of different sequences and in different conformations are shown in Figure 2b. The average number of sodium ions ranged from ~2.6 to ~6.6 and is strongly sequence dependent. In several sequences, (GT)_6_, C_12_, (TG)_3_(GT)_3_, ring conformations had larger number of sodium ions nearby on average than other conformations.

Representative structures of two sequences that had a strong propensity (C_12_) and no propensity (T_12_) for ring formation were examined by means of contact maps, shown in Fig. 3. The opacity of color in contact maps represents the probability of nucleic acid pairs being found within 4.5 Å of each other. Figure 3a shows contact maps of structural ensembles corresponding to ring, right-handed and left-handed helix conformations of C_12_. For all three conformations, each nucleotide has four highly probable neighbors: two nucleotides that are covalently linked to it, and two nucleotides that are its second neighbors ((n, n+2) interactions). The second-neighbor interactions are prominent due to compactness of ring and helical conformations. In ring conformations, the first and the last nucleotides are observed to interact while clasping the ring. In helical conformations, more nucleotides at 3’- and 5’-ends tend to interact.

**Figure 3.**
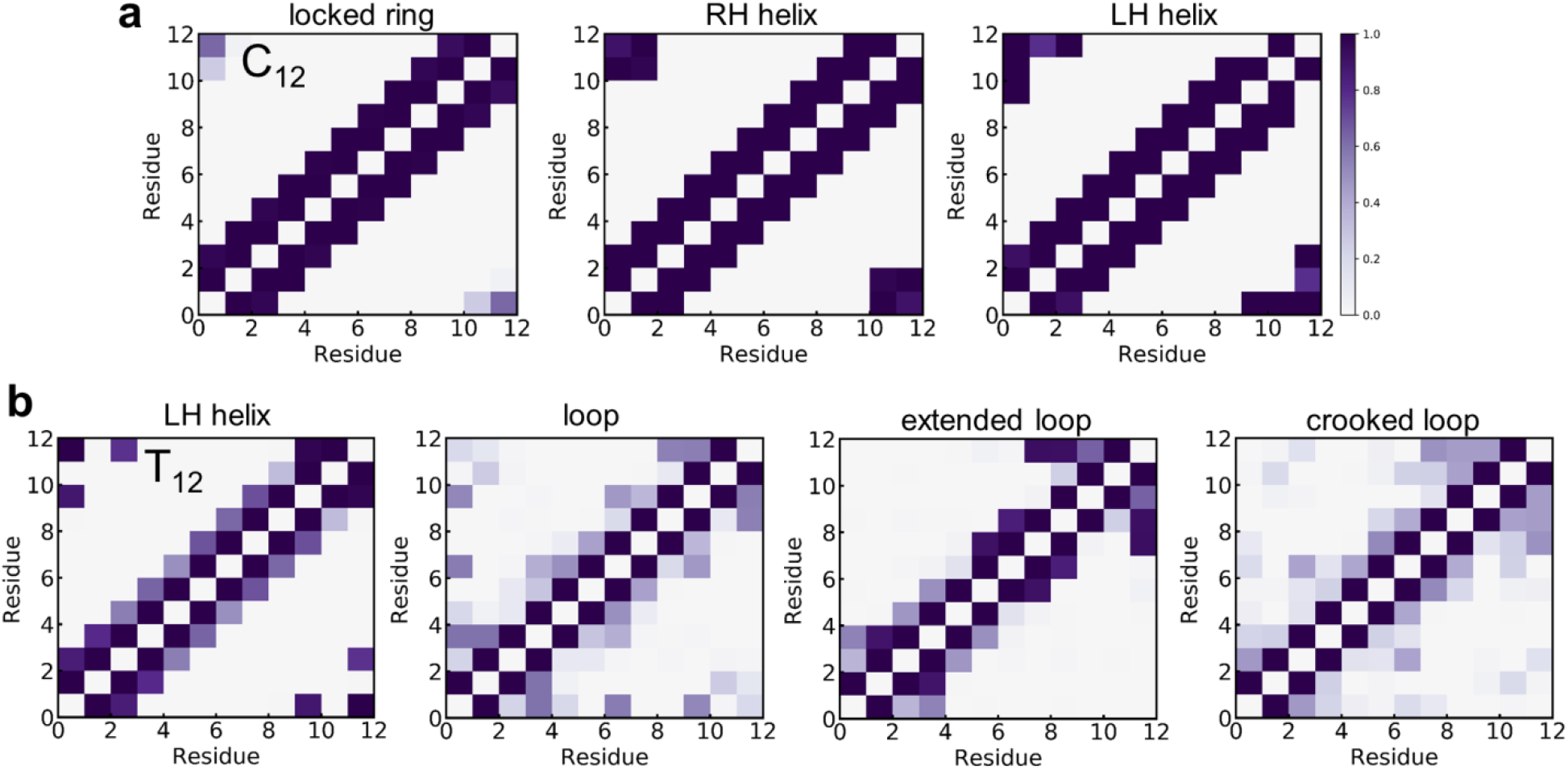
Contact maps of C_12_ and T_12_ molecules, where the structures are separated according to their free energy minimum basins. The structural ensembles of ssDNA molecules on (9,4) SWNTs were extracted from REMD simulations at 300 K. Abbreviations LH and RH refer to left-handed and right-handed.

Figure 3b shows contact maps of structural ensembles corresponding to left-handed helix and three different loop conformations of T_12_. Contact maps of T_12_ exhibits very different features from C_12_ maps. In T_12_, each nucleotide primarily interacts with its nearest covalently linked neighbors, and exhibits much weaker interactions with second neighbors ((n, n+2)). In T_12_, there are also more interactions of nucleotides with their non-immediate neighbors.

Interestingly, there is no correlation between the ssDNA composition (including purine, C-nt, T-nt, and G-nt fractions in ssDNA) and the ring formation propensity, as shown in Figure S4. Also, we examined if molecules in conformations in ring basins have contact areas with SWNTs that are distinct from conformations in other basins (Fig. S5). While contact area of C_12_ ring conformations was higher by ~100 Å^2^ compared to other conformations, (GT)_6_ ring conformations exhibited contact areas smaller by ~ 60 Å^2^ than the left-handed helix conformations. Therefore, contact area of ssDNA with SWNT did not seem to be correlated with the ring conformation.

### 2.4. Free energy landscapes at low ionic strength

Conformations of DNAs on SWNTs are predicted to have a pronounced dependence on the ionic strength^26^. Therefore, we examined how the free energy landscape of two representative sequences, C_12_ and T_12_, were affected by lowering the ionic strength. The free energy landscapes for these sequences at 150 mM and 1 mM NaCl solutions are shown in Figs. 2b-c and 4, respectively. We found that the free energy minima are in roughly the same parts of the reaction coordinate space for both ionic solutions. However, relative populations of these minima are shifted. For C_12_, minima 1 (ring) become less populated, minima 2 (right handed helix) become more populated, and minima 3 (left handed helix) become shallower and broader, likely due to more pronounced electrostatic repulsion between phosphate groups. For T_12_, locations of free energy minima are also similar for both ionic solutions. However, in 1 mM NaCl, minima were less separated by free energy barriers (transitions between different minima are more likely). In addition, other smaller local minima observed in 150 mM NaCl become more pronounced in 1 mM NaCl. Since free energy minima are located in roughly the same parts of the reaction coordinate space for both ionic solutions, our results seem to indicate that free energy landscapes for short ssDNAs are more sequence-dependent than solution ionic strength dependent. However, other intermolecular (inter-ssDNA) phenomena may emerge in low NaCl concentration solutions, which were not captured in the simulations performed.

**Figure 4.**
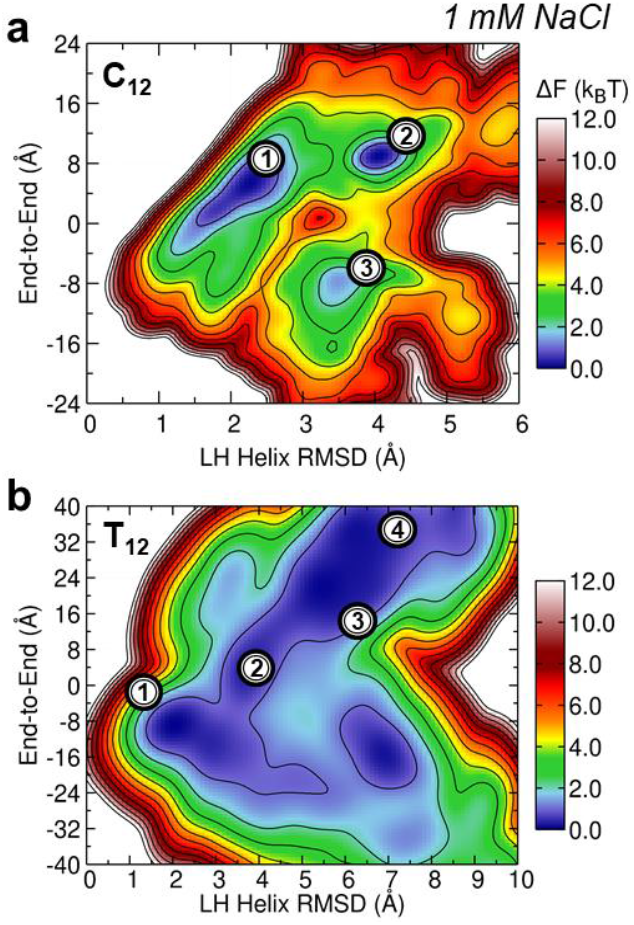
Free energy landscapes of a. C_12_ and b. T_12_ ssDNAs on (9,4) SWNTs at 300 K, in 1 mM NaCl aqueous solutions. The labels indicating free energy minima and their representative structures correspond to the labels shown in Fig. 1. Free energy landscapes for the same sequences in 150 mM NaCl aqueous solutions are shown in Fig. 1.

### 2.5. Kinetic stability of short oligonucleotides on SWNTs in experiments

To explore the relationship between sequence chemistry and kinetic stability, we performed surfactant displacement experiments on twelve screened sequences. Previous studies have shown that high affinity surfactants like sodium cholate (SC) can displace SWNT surface-adsorbed ssDNA^39–41^. We leveraged SC affinity for SWNT surface to obtain an experimental readout for the relationship between sequence chemistry and kinetic stability of the adsorbed species. Towards this end, we added SC to ssDNA-SWNT suspensions to displace the oligonucleotides adsorbed to the SWNT surface. All sequences display a blue-shift of center peak wavelengths, where each peak corresponds to different chiralities, suggesting displacement of the surface-adsorbed oligonucleotides by SC. Figures 5a and 5b describe the time-resolved solvatochromic shift of the (9,4) peak monitored over 300 seconds. Upon SC addition, the (9,4) peak shifts towards 1116 nm, which is the (9, 4) peak of SC-suspended SWNT, suggesting that SC has displaced most of the surface-adsorbed oligonucleotides. We identified four sequences that displayed minimal wavelength shift, exhibiting a change of approximately 1 nm over the time period monitored. We hypothesize that the extent of the blue-shift reflects the degree of displacement by SC and eventual surfactant surface coverage, and the speed with which the shift occurs indicates the ease of displacement of the oligonucleotide.

**Figure 5.**
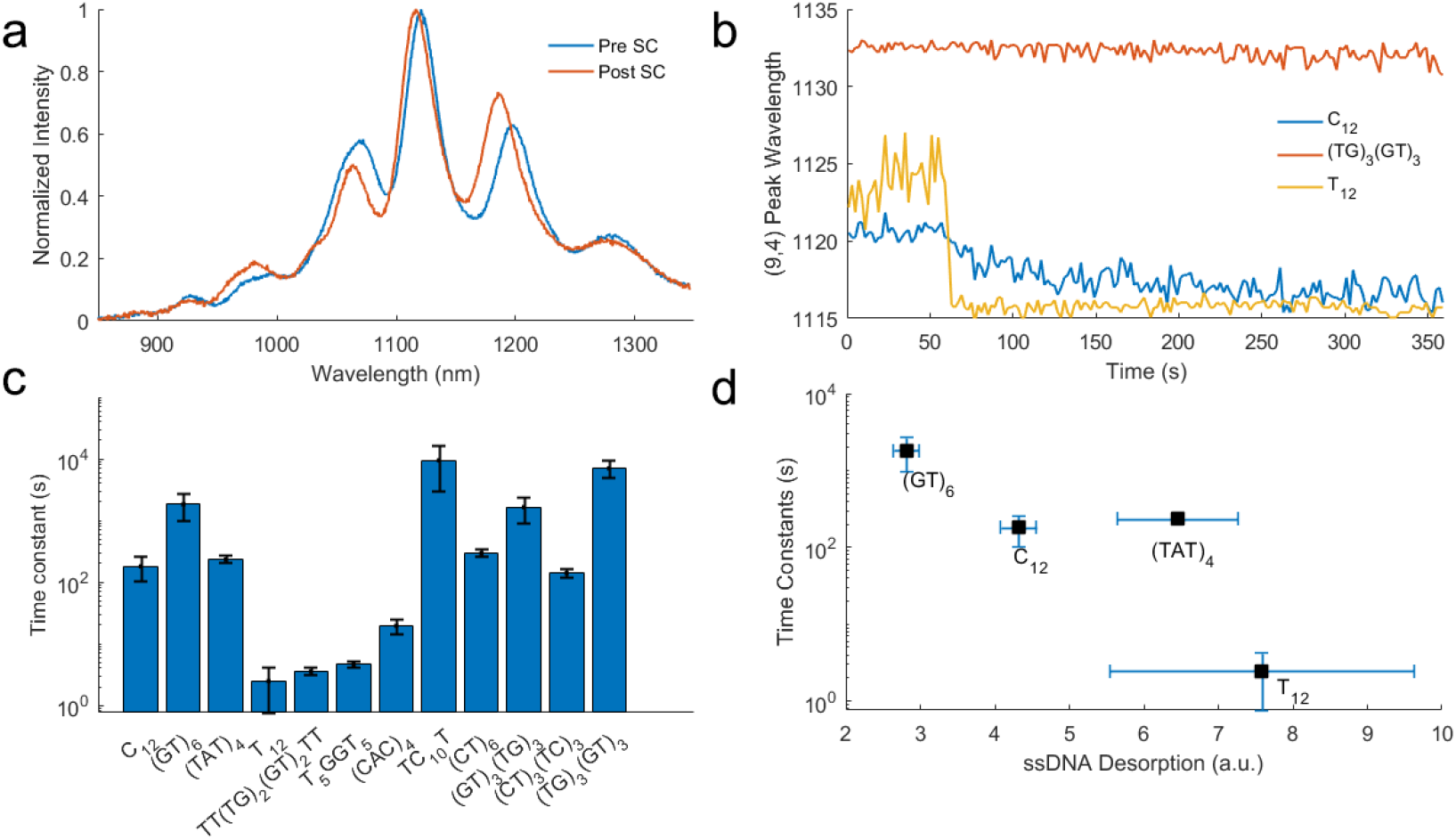
SC induced solvatochromatic shift is dependent on ssDNA sequence. (a) Addition of 0.1% SC induces a solvatochromic shift in (C_12_)-SWNT fluorescence. (b) Time resolved fluorescence measurements of (C_12_)-SWNT, (TG)_3_(GT)_3_-SWNT, and (T_12_)-SWNT upon addition of SC as indicated by arrow. The degree of solvatochromic shift is sequence dependent. (c) Time constants of solvatochromic shift is representative of how much time is needed for ssDNA to be displaced from the surface by surfactant. (d) Larger solvatochromic shift time constants are correlated to lower degrees of ssDNA desorption.

To quantitatively analyze the kinetics of the process, we assumed the temporal dynamics follows a first order kinetic model and fit the experimental data using least squares regression, from which decay time constants were determined (Figure 5c). The relative magnitude of the time constants are consistent with the speed of blue-shift we observe in the raw data, with larger time constants correlated with a slower blue-shift. Four sequences, (GT)_6_, TC_10_T, (GT)_3_(TG)_3_, and (TG)_3_(GT)_3_, were found to have time constants at least an order of magnitude larger than the rest. These four sequences also exhibited minimal blue-shifting (Figure S6). We hypothesize that the relative value of the time constants and extent of blue-shift reflects the relative kinetic stability of the oligonucleotide sequences on the SWNT surface, with a higher time constant representing a more robust stability of the oligonucleotide to adsorption on the SWNT.

We validated the relationship between ssDNA stability and ssDNA desorption from SWNT by performing a SWNT corona ligand tracking assay^28^. We hypothesize that ssDNA sequences that show shorter SC-induced time constants are likelier to desorb from the SWNT surface altogether. To determine the degree of SC-induced desorption of ssDNA, we tracked the ssDNA proximity to the SWNT with fluorophore quenching. We labeled four sequences of ssDNA, (GT)_6_, C_12_, (TAT)_4_, and T_12_, with a Cy5 fluorophore (ex/em 664/668 nm) on the 3’ end. The Cy5 fluorophore is initially quenched when ssDNA is on the SWNT surface, and the fluorescence increases in intensity upon ssDNA desorption from the SWNT^28^. SC was added to ssDNA-Cy5-SWNT in a well plate and the Cy5 fluorescence was measured over time to monitor the degree of ssDNA desorption. All sequences showed an increase in fluorescence over time (Figure S7), consistent with the prediction that upon the addition of SC, ssDNA is displaced from SWNT surface. Out of the four sequences tested, T_12_ displayed the largest fluorescence fold increase and (GT)_6_ the smallest, suggesting that T_12_ had the highest degree of desorption. Overall, we find that a larger SWNT fluorescence increase correlates with a smaller solvatochromic shift time constant (Figure 5d), suggesting that solvatochromic shift is a good indicator of the degree of ssDNA desorption from the SWNT. A larger time constant is associated with slower desorption, which agrees with the earlier hypothesis that the speed and extent of solvatochromic shift correlates with ssDNA kinetic stability. As a control, when SC is added to a solution of dye-labeled ssDNA, only marginal increases in fluorescence were observed, likely driven by changes in solvent dielectric constant (Table S4). Therefore, this corona ligand tracking assay is consistent with data collected from SWNT fluorescence alone and the results in sum offer an experimental insight into kinetic stability of adsorbed species.

Next, we examine the influence of oligonucleotide binding affinity and its favored conformation on its kinetic stability on SWNT, as measured by experiments. The processes of oligonucleotide binding and conformational changes on SWNTs, schematically shown in Fig. 6c, are driven by its search for the free energy minimum state on SWNT. We hypothesized that both the binding affinity and the nature of the favored conformation may affect the ease of displacement of oligonucleotides on SWNT surface by surfactants and therefore its kinetic stability. To estimate the binding affinity of each sequence to SWNT, a simple model was prepared, based on summation of single nucleotide binding free energies to SWNT for all the nucleotides within single strands. The single nucleotide binding free energies are based on the values reported in Ref. 34, estimated for a system containing (5,5) CNT (A: −11.3 kcal/mol, G: −13.145 kcal/mol, C: −6.92 kcal/mol, T: −11.76 kcal/mol). The estimated binding affinities of several sequences are reported in Fig. S8.

**Figure 6.**
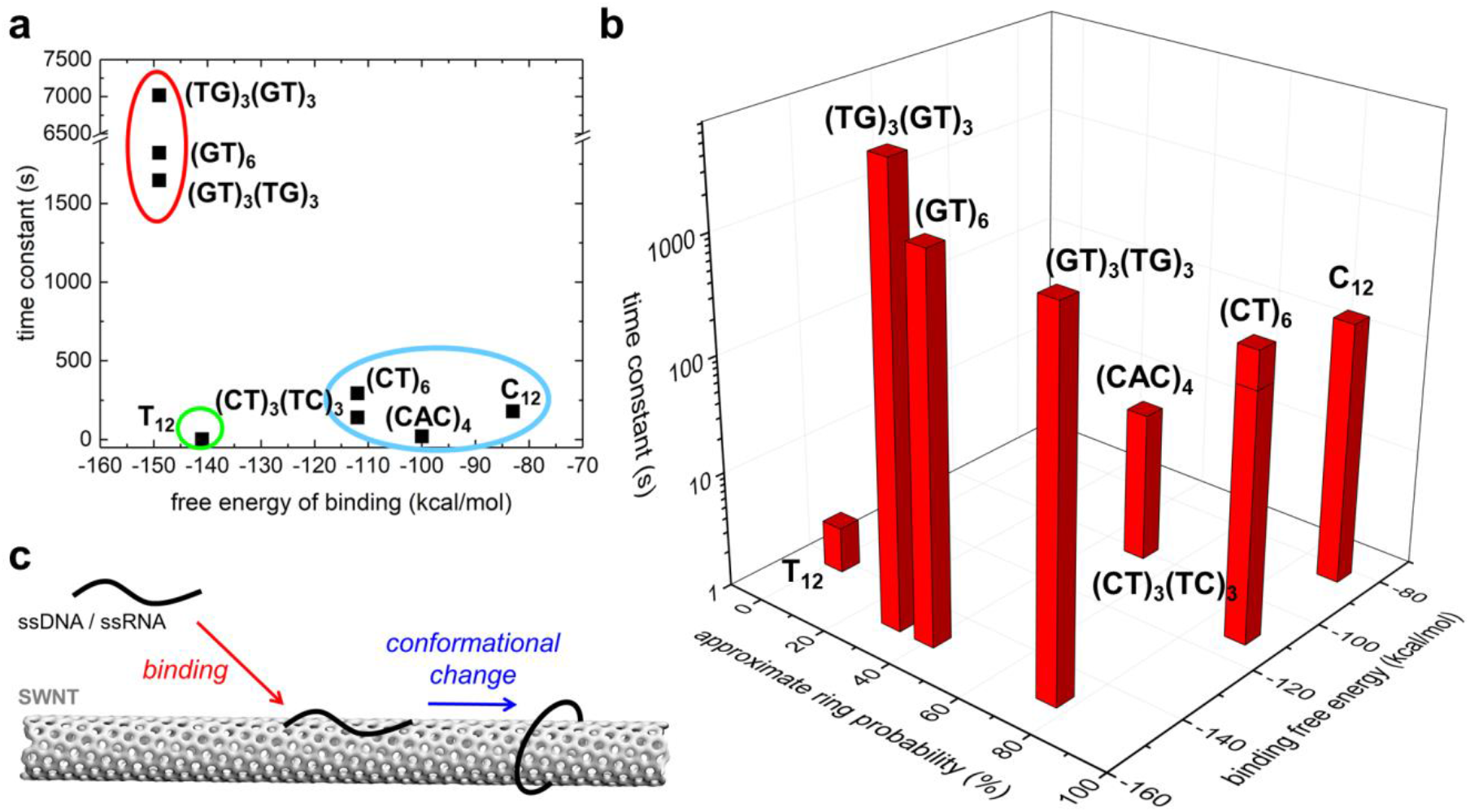
Correlations between computational and experimental data. a. Experimentally determined time constants of select ssDNA sequences versus their estimated free energies of binding to SWNT (Fig. S8). b. Experimentally determined time constant of select ssDNA sequences versus the estimated free energy of binding (Fig. S8) and their approximate probability to form ring conformation. c. A scheme of the processes associated with the adsorption of oligonucleotides to (9,4) SWNTs: initial binding of oligonucleotides, followed by conformational changes between favorable confirmations, revealed in the free energy landscapes. For (TG)_3_(GT)_3_, structures from ring and compact left-handed helix (partly) basins contribute to the compact structures approximated along the approximate ring probability axis.

Figure 6a shows the plot of the experimentally determined time constants of selected ssDNA sequences, extracted from Fig. 5, versus their estimated free energies of binding (adsorption) to SWNT, ΔG_ads_, extracted from Fig. S8. Clear correlations are observed between ΔG_ads_ and the time constant, i.e. the kinetic stability, for all of the sequences shown, except for an outlier T_12_ sequence. The observed correlations confirm that the ease of displacement of oligonucleotides on SWNTs by surfactants depends on how strongly the oligonucleotides bind to SWNT, as measured by ΔG_ads_.

To examine if the nature of the favored conformation of the oligonucleotide also affects its kinetic stability, the relationship between ΔG_bind_, favored conformation of the sequence, and the kinetic stability of selected sequences is plotted in Fig. 6b. The favored conformations are classified as either ring/helix, in which the oligonucleotides are “ordered” via strong (n,n+2)-nucleotide interactions, or loop conformations, in which oligonucleotides are more disordered. Again, Fig. 6b confirms that the binding affinity of the sequence to CNT has a clear correlation to its kinetic stability. The favored conformation then has a secondary influence on kinetic stability: the “ordered” conformations are somewhat correlated with slightly increased kinetic stability of sequence, as seen, for example, for sequences with weaker affinity to CNT (sequences with a large fraction of C-nucleotides, ΔG_bind_ > −120kcal/mol). We note that intermolecular interactions are not accounted for in the analysis, but could play an important role and lead to further modulation or a larger change in kinetic stability.

## 3. CONCLUSION

In conclusion, we expanded the library of short oligonucleotide sequences that formed ring-like structural motifs when adsorbed on SWNTs. Based on this work, we conclude that ring structural motifs on SWNTs emerge as a consequence of the oligonucleotide’s ability to form intramolecular hydrogen bonds, in particular between (n, n+2) nucleobases and between bases at the 3’ and 5’ ends of the polymer and is favored in sequences that are best able to accommodate sharp bends in their phospho-diester backbone. The propensity for ring formation is therefore likely an emergent property driven by the chemistry of the constituent bases and base stacking interaction with the SWNT surface. Our work however did not identify particular chemical trends (for example, purine vs. pyrimidine content) that reliably correlated with the formation of rings. Experimentally, we explored kinetic stability of the surface adsorbed polymers using a surfactant displacement assay. We identified that computed estimates of ΔG_ads_ are the primary correlates of experimentally measured stability. However, ring motifs contributed positively to kinetic stability. In sequences with unfavorable free energy of adsorption (more positive ΔG_ads_), ring structures contributed positively to stability. On the other hand, sequences that did not form rings but had favorable free energy of adsorption (more negative ΔG_ads_) were kinetically stable. Our work identified computationally determined parameters and their experimental correlates to better elucidate the structure-function relationship in oligonucleotide-SWNT hybrid systems.

## 4. METHODS

To understand the relationships between conformations, binding affinity and kinetic stability of short oligonucleotides on SWNTs, MD simulations and experiments of short ssDNA and ssRNA polymers on (9,4) SWNTs were performed.

### 4.1. Molecular dynamics simulations

Atomistic simulations were conducted to investigate conjugates of (9,4) SWNTs with 10-, 11- or 12-nucleotide ssDNAs and ssRNAs of different sequences, listed in Table S2. The same procedure was used to build the initial structures of conjugates of (9,4) SWNT with the nucleic acids. First, (9,4) SWNT segments, 39 Å in length, were built with VMD software^42^. The initial configurations of ssDNA and ssRNA molecules were built in Material Studio with nucleotides arranged to form helical conformations with radii several Ångstroms wider than the radius of the (9,4) SWNT. The helical DNAs were positioned to wrap SWNTs, with ssDNA bases not pre-adsorbed to the SWNTs surfaces. The prepared ssDNA-SWNT conjugates were solvated with TIP3P water and neutralized with 0.1 M NaCl with *solvate* and *ionize* VMD plugins, respectively^42^. The final systems contained approximately 6,300 atoms.

The systems were described with CHARMM36 parameters^43, 44^. MD simulations were performed with NAMD2.12 package^45^. All simulations were conducted with Langevin dynamics (Langevin constant Lang = 1.0 ps^−1^) in the NpT ensemble, where temperature and pressure remained constant at 310 K and 1 bar, respectively. The particle-mesh Ewald (PME) method was used to calculate Coulomb interaction energies, with periodic boundary conditions applied in all directions^46^. The time step was set to 2.0 fs. The evaluation of long range van der Waals and Coulombic interactions was performed every 1 and 2 time steps, respectively. After 1,000 steps of minimization, solvent molecules were equilibrated for 2 ns around the DNA and SWNTs, which were restrained using harmonic forces with a spring constant of 1 kcal/(molÅ^2^). Next, the systems were equilibrated in 80 ns production MD runs, with restraints applied only on the edge SWNT atoms.

### 4.2. Free energy calculations

The free energy landscapes (Figs. 1, 4, S2, S3) were obtained through replica exchange MD (REMD) simulation of ssDNA/ssRNA-SWNT systems. All systems were solvated in 3.63 × 3. 63 × 4.92 nm^3^ box, containing approximately 6,605 atoms. The box contained approximately 1,881 water molecules, modeled using the TIP3P model. In addition to Na^+^ counterions neutralizing the system, 45 Na^+^ and Cl^−^ ions were included to match the physiological salt concentration in the experimental system. Periodic boundary conditions were imposed in all dimensions, and PME method was used to calculate long-range electrostatics. Additionally, both ends of SWNT were in contact with their periodic images. Energy minimization and 100 ps of heating (NVT) were performed to reach the starting temperature of 310 K. To perform REMD simulations in NVT ensemble, 54 replicas and a 290-727.4 K temperature range were chosen to maintain exchange acceptance ratios around 25% with 2 ps exchange time. The total REMD simulation times are summarized in Table S1. The simulation time step was 2 fs and trajectories were extracted every 2 ps.

The first 10,000 (20 ns) configurations of room temperature replica were ignored, and the rest of the configurations were analyzed to obtain the free energy landscape. To generate the free energy landscapes shown in Figs. 1, 4, S2, and S3, two independent order parameters of ssDNA structures were calculated from the obtained system configurations: 1) end-to-end distance of DNA molecule (the z-distance between centers of mass of the first nucleotide and last nucleotide; z coordinate aligns with the long axis of the CNT); and 2) root mean square deviation (RMSD) of phosphorous atoms of the DNA backbone, compared to the configuration these atoms have in the ideal left-handed helix of ssDNA/ssRNA wrapping SWNT.

The probability distribution function (P(x,y)) of the two order parameters described above were calculated and combined to generate free energy (F(x; y)) according to the formula:

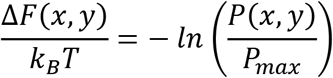

In the above formula, P_max_ is the maximum value of P(x, y). All the free energy landscapes shown were obtained at 300 K by calculating the two-dimensional free energy landscape according to the above formula, where x and y represent the two selected reaction coordinates described above (the end-to-end distance of DNA molecule and RMSD of P-atoms).

### 4.3. Data Analysis

#### Contact areas

Contact areas between nucleic acids and SWNT were calculated based on the following equation:

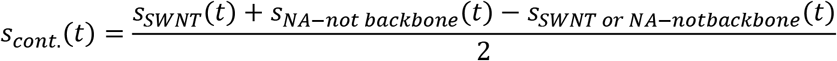

where *S*_*SWNT*_(*t*) and *S*_*NA−not backbone*_(*t*) represent solvent accessible surface areas (SASA) of SWNT and nucleic acid atoms except nucleic-acid backbone atoms at time t, respectively. *S*_*SWNT*_(*t*) or *S*_*NA−notbackbone*_(*t*) denotes SASA of a group of atoms including both SWNT and nucleic acid atoms except backbone atoms. The contact areas were calculated separately for all the structures that populate local free energy minima in free energy landscapes obtained at 300 K. The calculations were performed with SASA VMD plugin^42^. In SASA calculations, van der Waals radius of atoms was set to be 1.4 Å.

#### Contact maps

Contact maps of DNA residues wrapping SWNT were calculated with MDtraj^47^. The cut-off distance of 0.45 nm was selected for counting the number of residues that are in contact.

#### The probability of observing ring conformations

To determine the probability of observing ring conformations in short MD simulation trajectories, we evaluated for all the sequences and for all the time frames 1) the end-to-end distance of DNA molecule (the z-distance between centers of mass of the first nucleotide and last nucleotide; z coordinate aligns with the long axis of the CNT) and 2) the root mean square deviation (RMSD) of phosphorous atoms of the DNA backbone, compared to the configuration these atoms have in the ideal left-handed helix of ssDNA/ssRNA wrapping SWNT (a single reference structure). Three dimensional scatter plots of the end-to-end distances and the RMSDs for each time frame were prepared and examined. In the analysis, the clusters of points corresponding to ring conformations were defined to have end-to-end distances in the range (−4 Å to 4 Å) and RMSD > 2.5 Å. The ring conformations were either transient (the points in the scatter plots did not form a cluster) or stable (the points in the scatter plot formed a cluster). For each sequence, the number of points corresponding to ring conformations was summed and divided by the total number of points comprising each trajectory (usually 4,000), providing the fractions of time that the molecules assumed ring conformations in the calculated trajectories.

### 4.4 Experimental studies

ssDNA oligonucleotides were purchased from Integrated DNA Technologies (Standard Desalting) and HiPCo SWNT were purchased from NanoIntegris (Batch #HR27-104). 1 mg ssDNA was mixed with 4 mg SWNT in 1 mL of 0.1 M NaCl solution. The solution was bath sonicated (Branson Ultrasonic 1800) for 15 minutes and probe-tip sonicated with 3 mm tip (Cole Parmer Ultrasonic Processor) for 10 minutes at 6 W in an ice bath. The sonicated solution was allowed to equilibrate at room temperature for 30 minutes. Unsuspended SWNT and other amorphous carbon material was precipitated by centrifuging the sonicated solution at 16,100 g for 60 minutes. The supernatant was recovered and the concentration of the colloidally stable ssDNA-SWNT was determined with UV-Vis measurements (ThermoFisher NanoDrop One) using an extinction coefficient of 0.036 (mg/L)^−1^ cm^−1^ at 632 nm^26^. For the preparation of ssDNA-Cy5-SWNT, 20 μM ssDNA with 3’ labeled cyanine-5 (Cy5) was mixed with 1 mg SWNT in 0.01M phosphate-buffered saline (PBS) and the previous suspension protocol was employed with the following changes: the solution was not bath sonicated and the centrifugation time was decreased to 30 minutes. The concentration of ssDNA-Cy5-SWNT was determined with UV-Vis measurements with an extinction coefficient of 0.02554 (mg/L)^−1^ cm^−1^ at 910 nm^48^.

A 5 wt.% sodium cholate (SC) solution was prepared in Milli-Q water. ssDNA-SWNT solutions were diluted to 5 mg/L in 0.1 M NaCl solution and 200 μL aliquots were added to each well of a 96-well well plate. Fluorescence emission spectra from each well was determined with a 10x objective on an inverted Zeiss microscope (Axio Observer.D1) connected to a Princeton Instruments spectrometer (SCT 320) and coupled to a liquid nitrogen-cooled Princeton Instruments InGaAs detector (PyLoN-IR). A 721 nm laser (OptoEngine LLC) produced 80 mW laser power at the back focal plane of the objective for excitation. Spectra were recorded with 1 second exposure time. To obtain temporally resolved evolution of fluorescence shift caused by SC, we first monitored emission from each well for 60 seconds to obtain baseline spectra. To record the fluorescence emission wavelength shift, we added 4 μL of the 5 wt.% SC solution to a well as marked by a black arrow in Figure 5b, and SC was allowed to diffuse passively through the well. After SC addition, fluorescence emission spectra was collected for an additional 300 seconds. All measurements were background corrected with blank 0.1 M NaCl solution.

40 μL of ssDNA-Cy5-SWNT in 0.1 M PBS and 10 μL of SC solution were added to a 96-well PCR plate (Bio-Rad) for a final concentration of 5 mg/L ssDNA-Cy5-SWNT and 0.1 wt.% SC. For control measurements, either ssDNA-Cy5-SWNT was substituted with ssDNA-Cy5 at a final concentration 0.20 μM or SC was substituted with 0.1 M PBS. The plate was covered with an optically transparent adhesive seal (Bio-Rad). Cy5 fluorescence was measured with a Bio-Rad CFX96 Real Time qPCR System, scanning all manufacturer set color channels (FAM, HEX, Texas Red, Cy5, Quasar 705) every 30 seconds at 22.5 °C (no lid heating) for a total of 60 minutes. Time-resolved fluorescence was analyzed without background correction^28^. Fluorescence fold increase was calculated by dividing the Cy5 intensity of ssDNA-Cy5-SWNT with SC added by the Cy5 intensity of ssDNA-Cy5-SWNT with PBS.

### 4.5 TEM Imaging

For TEM imaging of (GT)_6_-SWNT and (GT)_15_-SWNT samples, a FEI ThemIS TEM with 60kV acceleration voltage was used. An aliquot of 5μL of (GT)_6_-SWNT or (GT)_15_-SWNT at a concentration of 10mg/L was drop casted onto holey carbon film coated Cu TEM grids, allowed to adsorb for 30 minutes, and the surface was rinsed. The TEM grid surface was treated with a glow plasma discharger for hydrophilic surfaces. Samples were used directly for TEM imaging without staining.

## 5. ACKNOWLEDGMENTS

The authors gratefully acknowledge computer time provided by the Texas Advanced Computing Center (TACC). This research is part of the Blue Waters sustained-petascale computing project, which is supported by the National Science Foundation (Awards OCI0725070 and ACI-1238993) and the state of Illinois. M.P.L. acknowledges support of a Burroughs Wellcome Fund Career Award at the Scientific Interface (CASI), a Stanley Fahn PDF Junior Faculty Grant with Award # PF‐JFA‐1760, a Beckman Foundation Young Investigator Award, a DARPA Young Faculty Award, an FFAR New Innovator Award, and a USDA award. M. P. L. is a Chan Zuckerberg Biohub Investigator and an Innovative Genomics Institute Investigator.

