## Supplemental Information for "Binding affinity and conformational preferences influence kinetic stability of short oligonucleotides on carbon nanotubes"

**Table S1.** Detailed information on REMD systems.

| Systems | Total number of atoms | Simulation time (ns) | Number of configurations (ns) |
|---|---|---|---|
| $(GT)_6$ | 6605 | 270 | 80000 (160 ns) |
| $(C)_{12}$ | 6614 | 220 | 99000 (180 ns) |
| $(T)_{12}$ | 6601 | 124 | 52000 (104 ns) |
| $(CT)_6$ | 6679 | 220 | 99000 (198 ns) |
| $(AC)_6$ | 6614 | 200 | 90000 (180 ns) |
| $(CU)_6$ | 6603 | 220 | 90000 (180 ns) |
| $(CU)_5UU$ | 6609 | 220 | 90000 (180 ns) |

**Table S2.** Screening short single stranded nucleic acids for ring conformation probability on (9,4) SWNTs. For RNA molecules, most of the observed rings are significantly tilted with respect to the SWNT axis.

| | NA sequences - (9,4) CNT | Number of nucleotides | Ring stability |
|---|---|---|---|
| 1 | $TT(TG)_2(GT)_2TT$ | 12 | 0.014 |
| 2 | $TT(GT)_2(TG)_2T$ | 11 | 0.254 |
| 3 | $TT(TG)_2(GT)_2T$ | 11 | 0.023 |
| 4 | $TT(GT)_2(TG)_2$ | 10 | 0.288 |
| 5 | $TG(GT)_2(TG)_2$ | 10 | 0.41 |
| 6 | $(TGT)_3TG$ | 11 | 0.099 |
| 7 | $(T)_5GG(T)_5$ | 12 | 0 |
| 8 | $(T)_5GG(T)_4$ | 11 | 0.0915 |
| 9 | $(T)_4GG(T)_4$ | 10 | 0.1 |
| 10 | $(T)_{12}$ | 12 | 0 |
| 11 | $(T)_{12}$-1nM salt | 12 | 0 |
| 12 | $(ATTT)_3$ | 12 | 0.064 |
| 13 | $(TAT)_4$ | 12 | 0.1305 |
| 14 | $(CAC)_4$ | 12 | 0.189 |
| 15 | $(TAT)_3TA$ | 11 | 0.127 |
| 16 | $(TAT)_3TT$ | 11 | 0.2043 |
| 17 | $(TAT)_3T$ | 10 | 0.098 |
| 18 | $G(C)_{10}G$ | 12 | 0 |
| 19 | $T(C)_{10}T$ | 12 | 0.0025 |
| 20 | $T(C)_{10}$ | 11 | 0.124 |
| 21 | $G(C)_{10}$ | 11 | 0.1323 |
| 22 | $(C)_6(T)_6$ | 12 | 0.0048 |
| 23 | $T(C)_5(T)_6$ | 12 | 0 |
| 24 | $T(C)_5(T)_5$ | 11 | 0.168 |
| 25 | $(CT)_2(TC)_2(CT)_2$ | 12 | 0.0405 |
| 26 | $(GT)_3(TG)_3$ | 12 | 0 |
| 27 | $(TG)_3(GT)_3$ | 12 | 0.074 |
| 28 | $(CT)_3(TC)_3$ | 12 | 0.606 |
| 29 | $(C)_{12}$ | 12 | 0.16 |
| 30 | $(CT)_6$ | 12 | 0.07 |
| 31 | $(AC)_6$ | 12 | 0.54 |
| 32 | $(GT)_6$ | 12 | 0.275 |
| 33 | $(CU)_5C$ | 11 | 0.01 |
| 34 | $(CU)_5$ | 10 | 0.28 |
| 35 | $(CU)_6$ | 12 | 0.49 |
| 36 | $(CU)_5UU$ | 12 | 0.33 |

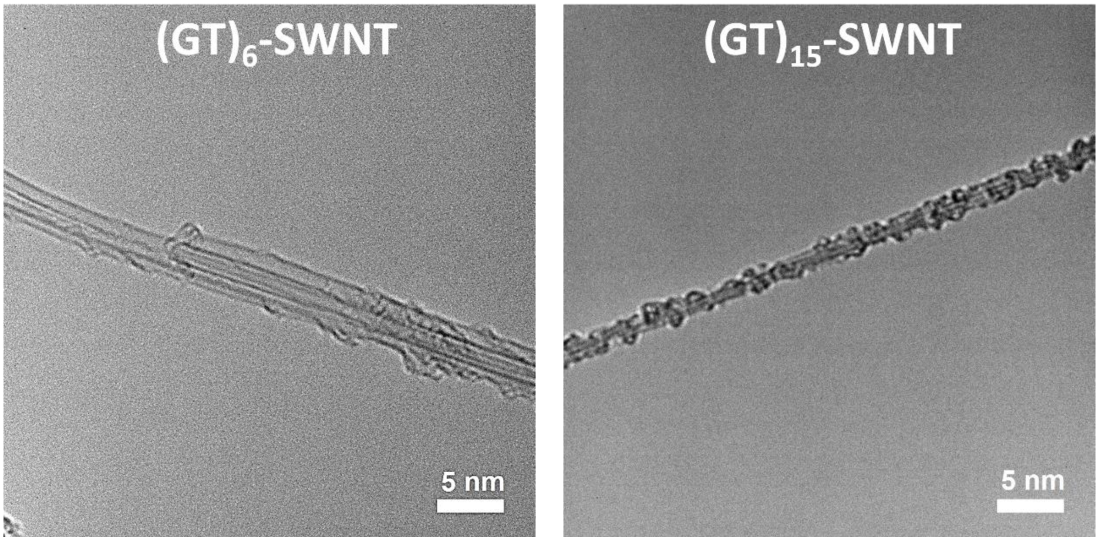


**Figure S1.** High-resolution TEM images of $(GT)_6$ and $(GT)_{15}$ polymers on CNTs.

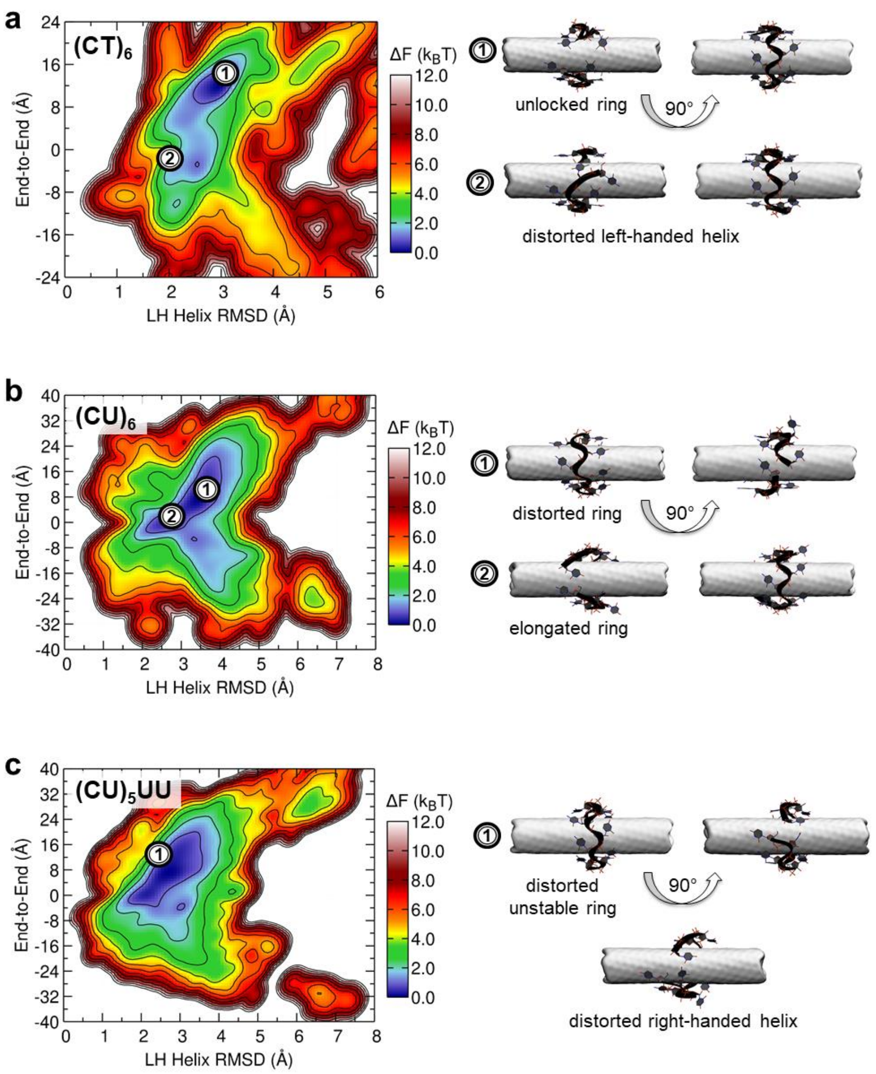


**Figure S2.** Free energy landscapes of 12-nt ssDNAs and ssRNAs on (9,4) SWNTs at 300 K. a. $(CT)_6$-SWNT (DNA). b. $(CU)_6$-SWNT (RNA). c. $(CU)_5UU$-SWNT (RNA).

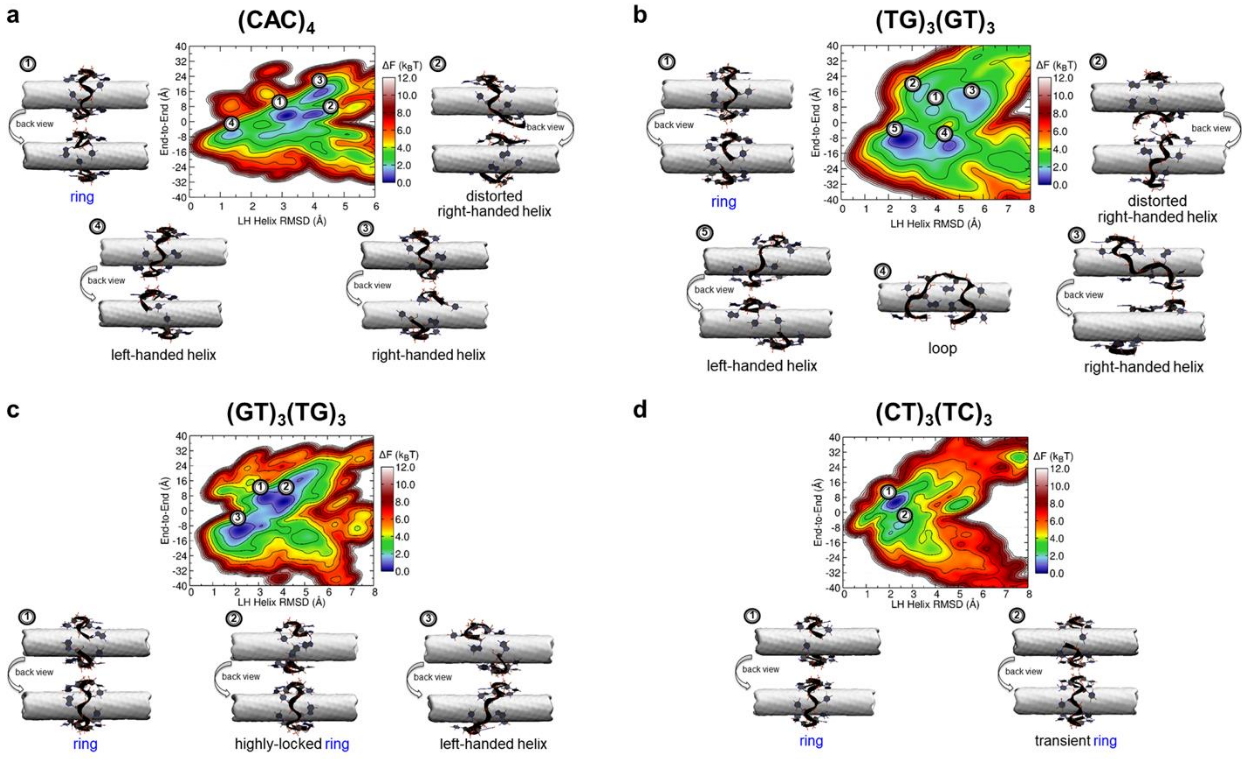


**Figure S3.** Additional free energy landscapes of 12-nt ssDNAs on (9,4) SWNTs at 300 K. a. $(CAC)_4$-SWNT. b. $(TG)_3(GT)_3$-SWNT. c. $(GT)_3(TG)_3$-SWNT. d. $(CT)_3(TC)_3$-SWNT.

**Table S3.** Average number of sodium ions within 4 Å of nucleic acids examined, averaged over all the structures in free energy minimum basins, determined from T-REMD simulations at 300 K. Abbreviations LH and RH refer to left-handed and right-handed.

| | | | | | |
|---|---|---|---|---|---|
| **$(GT)_6$** | ring | 6.13 | **$(CT)_3(TC)_3$** | ring | 3.94 |
| | LH helix | 5.33 | | transient ring | 3.83 |
| **$C_{12}$** | ring | 3.93 | **$(TG)_3(GT)_3$** | ring | 6.6 |
| | RH helix | 3.22 | | distorted RH helix | 4.83 |
| | LH helix | 2.62 | | RH helix | 4.75 |
| **$(CT)_6$** | unlocked ring | 3.58 | | loop | 4.77 |
| **$(AC)_6$** | extended ring | 4.74 | | LH helix | 5.55 |
| | ring | 4.6 | **$(GT)_3(TG)_3$** | ring | 6.14 |
| **$T_{12}$** | LH helix | 4.48 | | highly locked ring | 5.3 |
| | loop | 3.97 | | LH helix | 6.12 |
| | extended loop | 4.17 | | | |
| | crooked loop | 4.54 | | | |
| **$(CAC)_4$** | ring | 4.01 | | | |
| | distorted RH helix | 3.87 | | | |
| | RH helix | 4.31 | | | |

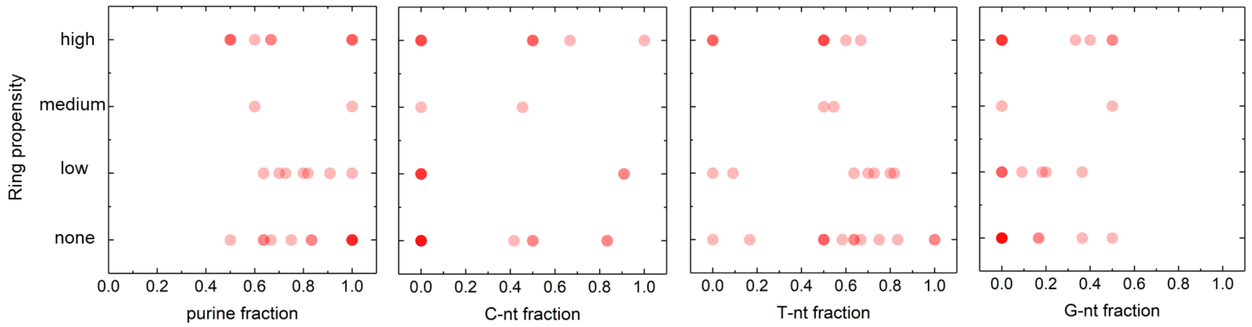


**Figure S4.** Ring propensity versus the ssDNA composition, including the purine, C-nt, T-nt, and G-nt fractions.

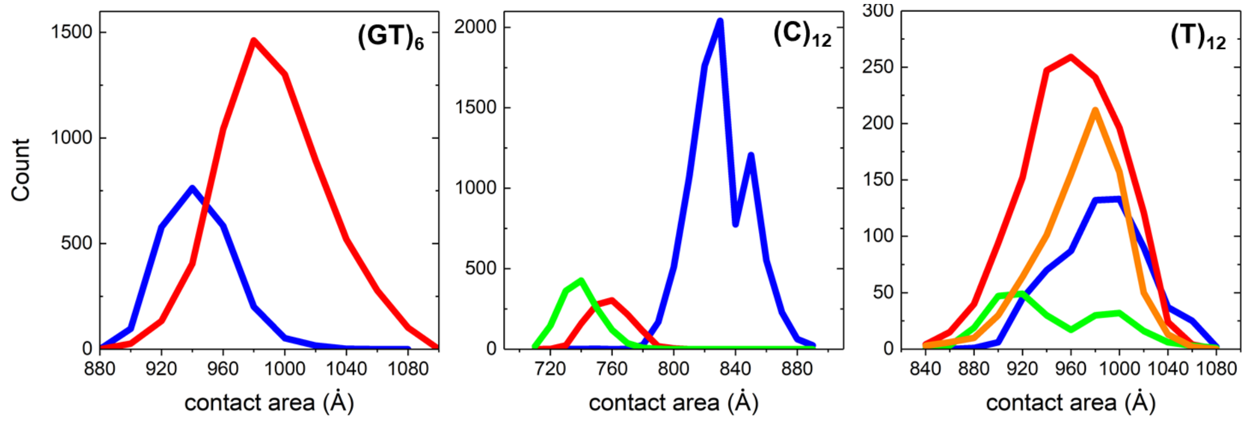


**Figure S5.** ssDNA-SWNT contact areas for $(GT)_6$-SWNT, $(C)_{12}$-SWNT, and $(T)_{12}$-SWNT ssDNA molecules on (9,4) SWNTs at 300 K.

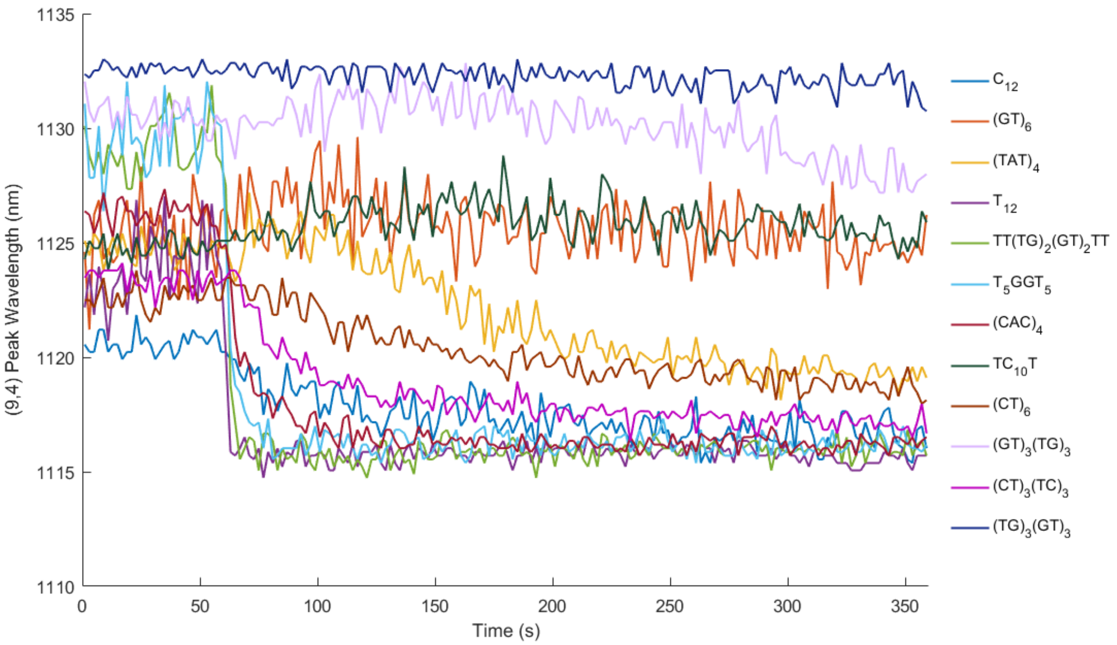


**Figure S6.** Time resolved fluorescence measurements of the (9,4) peak SWNT wavelength. SC was added to ssDNA-SWNT at 60 seconds as indicated by the black arrow. Sequences display blue-shift that approaches peak wavelength of SC-suspended SWNT.

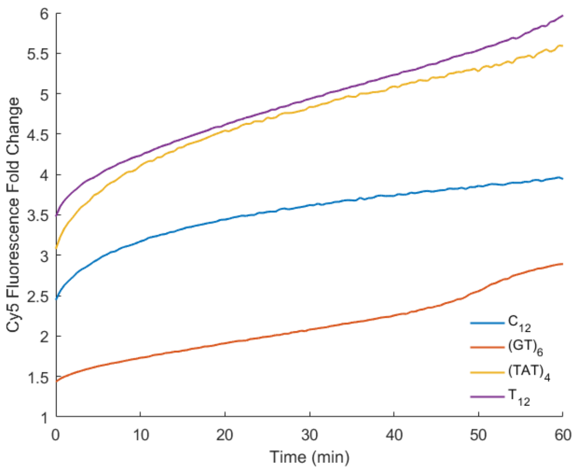


**Figure S7.** Time resolved fluorescence measurement of Cy5-labeled ssDNA in well plate. SC was added to ssDNA-Cy5-SWNT approximately 1 minute before the measurement was initiated. Cy5 fluorescence intensity increases after the addition of SC.

**Table S4.** Cy5 fluorescence fold change and solvatochromic shift time constant for four sequences after addition of SC. Larger fluorescence fold change is associated with a smaller solvatochromic shift time constant.

| Series | Cy5 fluorescence fold change | Solvatochromic shift time constant (s) |
|---|---|---|
| **$C_{12}$-Cy5-SWNT** | 4.32 ± 0.24 | 179.32 ± 77.02 |
| **$C_{12}$-Cy5** | 1.93 ± 0.56 | N/A |
| **$(GT)_6$-Cy5-SWNT** | 2.81 ± 0.17 | 1821.64 ± 851.40 |
| **$(GT)_6$-Cy5** | 1.53 ± 0.66 | N/A |
| **$(TAT)_4$-Cy5-SWNT** | 6.45 ± 0.81 | 233.07 ± 32.94 |
| **$(TAT)_4$-Cy5** | 1.91 ± 0.10 | N/A |
| **$T_{12}$-Cy5-SWNT** | 7.58 ± 2.04 | 2.47 ± 1.71 |
| **$T_{12}$-Cy5** | 2.14 ± 0.06 | N/A |

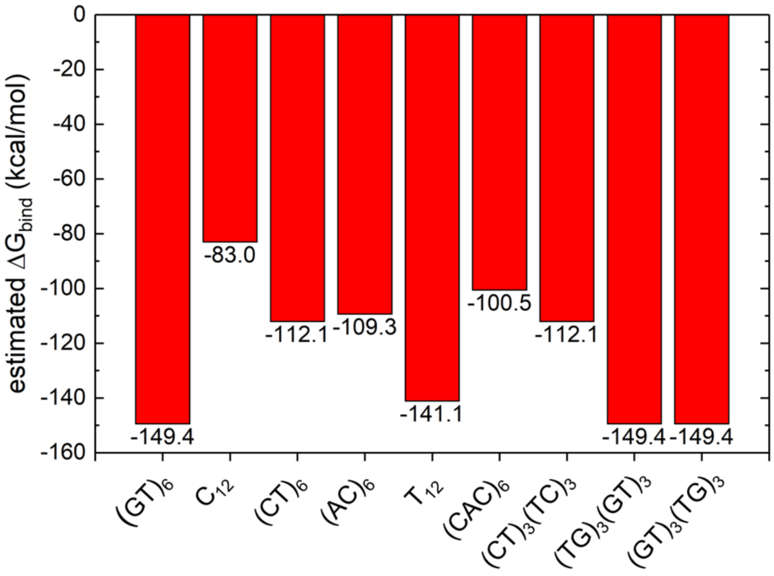


**Figure S8.** Estimated binding free energies of ssDNA strands to CNTs. The total estimated binding free energies of whole strands, reported on the y-axis, are based on the summation of single nucleotide binding free energies to CNT for all the nucleotides within single strands. The single nucleotide binding free energies are based on the values reported in Ref. 1, estimated for a system containing (5,5) CNT (A: -11.3 kcal/mol, G: -13.145 kcal/mol, C: -6.92 kcal/mol, T: -11.76 kcal/mol).

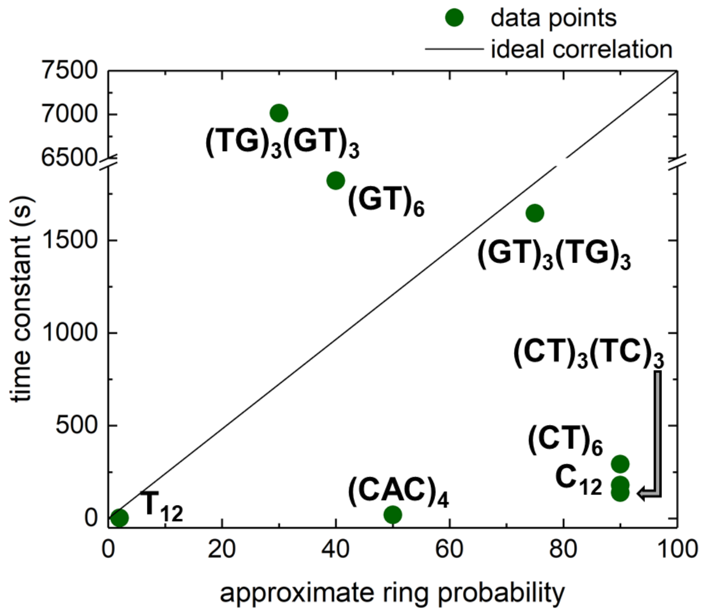


**Figure S9.** Correlations between computational and experimental data: experimentally determined time constants of select ssDNA sequences versus their approximate probability to form ring conformation. For $(TG)_3(GT)_3$, structures from ring and compact left-handed helix (partly) basins contribute to the compact structures approximated along the approximate ring probability axis.
